# Altered EEG resting-state large-scale brain network dynamics in euthymic bipolar disorder patients

**DOI:** 10.1101/668004

**Authors:** Alena Damborská, Camille Piguet, Jean-Michel Aubry, Alexandre G. Dayer, Christoph M. Michel, Cristina Berchio

## Abstract

**Background:** Neuroimaging studies provided evidence for disrupted resting-state functional brain network activity in bipolar disorder (BD). Electroencephalographic (EEG) studies found altered temporal characteristics of functional EEG microstates during depressive episode within different affective disorders. Here we investigated whether euthymic patients with BD show deviant resting-state large-scale brain network dynamics as reflected by altered temporal characteristics of EEG microstates.

**Methods:** We used high-density EEG to explore between-group differences in duration, coverage and occurrence of the resting-state functional EEG microstates in 17 euthymic adults with BD in on-medication state and 17 age- and gender-matched healthy controls. Two types of anxiety, state and trait, were assessed separately with scores ranging from 20 to 80.

**Results:** Microstate analysis revealed five microstates (A-E) in global clustering across all subjects. In patients compared to controls, we found increased occurrence and coverage of microstate A that did not significantly correlate with anxiety scores.

**Conclusion:** Our results provide neurophysiological evidence for altered large-scale brain network dynamics in BD patients and suggest the increased presence of A microstate to be an electrophysiological trait characteristic of BD.

## 1 Introduction

Bipolar disorder (BD) is a common and severe psychiatric disorder, with an important personal and societal burden (Cloutier et al., 2018; Eaton et al., 2012). The worldwide prevalence of BD is considered to range between 1% and 3% (Merikangas et al., 2011; Ferrari et al., 2016). BD patients are frequently misdiagnosed and often identified at late stages of disease progression, which can lead to inadequate treatment (Hirschfeld, 2007) and worse functional prognosis (Vieta et al., 2018). A better understanding of the underlying pathophysiology is needed to identify objective biomarkers of BD that would improve diagnostic and/or treatment stratification of patients.

Possible candidates for neurobiological biomarkers in BD could arise from the abnormalities of functional brain networks. Evidence from brain imaging studies consistently points to abnormalities in circuits implicated in emotion regulation and reactivity. Particularly, attenuated frontal and enhanced limbic activations are reported in BD patients (Chen et al., 2011; Houenou et al., 2011; Kupferschmidt and Zakzanis, 2011). Interestingly, regions implicated in the pathophysiology of the disease, such as the inferior frontal gyrus, the medial prefrontal cortex (mPFC), and the amygdala present altered activation patterns even in unaffected first-degree relatives of BD patients (Piguet et al., 2015), pointing towards brain alterations that could underlie disease vulnerability. Moreover, evidence from functional magnetic resonance imaging (fMRI) studies showed aberrant resting-state functional connectivity between frontal and meso-limbic areas in BD when compared to healthy controls (Vargas et al., 2013). A recently developed functional neuroanatomic model of BD suggests, more specifically, decreased connectivity between ventral prefrontal networks and limbic brain regions including the amygdala (Strakowski et al., 2012; Chase and Phillips, 2016). The functional connectivity abnormalities in BD in brain areas associated with emotion processing were shown to vary with mood state. A resting-state functional connectivity study of emotion regulation networks demonstrated that subgenual anterior cingulate cortex (sgACC)-amygdala coupling is critically affected during mood episodes, and that functional connectivity of sgACC plays a pivotal role in mood normalization through its interactions with the ventrolateral PFC and posterior cingulate cortex (Rey et al., 2016). Nevertheless, although different fMRI metrics allowed to report deviant patterns of large-scale networks and altered resting-state functional connectivity (Rey et al., 2016; Wang et al., 2016) in BD, the precise temporal dynamics of the functional brain networks at rest remain to be determined.

Large-scale neural networks dynamically and rapidly re-organize themselves to enable efficient functioning (de Pasquale et al., 2018; Bressler and Menon, 2010). Fast dynamics of the resting-state large-scale neural networks can be studied on sub-second temporal scales with EEG microstate analysis (Pascual-Marqui et al., 1995; Van de Ville et al., 2010; Michel and Koenig, 2018). EEG microstates are defined as short periods (60-120 ms) of quasi-stable electric potential scalp topography (Lehmann et al., 1987; Koenig et al., 2002). Therefore, microstate analysis can cluster the scalp’s topographies of the resting-state EEG activity into the set of a few microstate classes including the four canonical classes A-D (Michel and Koenig, 2018) and more recent additional ones (Custo et al., 2017; Bréchet et al., 2019). Since each microstate class topography reflects a coherent neuronal activity (Khanna et al., 2015; Michel and Koenig, 2018), the temporal characteristics, such as duration, occurrence and coverage, may be linked to the expression of spontaneous mental states and be representative of the contents of consciousness (Changeux and Michel, 2004; Lehmann, 1990). Numerous studies reported abnormalities in temporal properties of resting-state EEG microstates in neuropsychiatric disorders (for review see Khanna et al., 2015; Michel and Koenig, 2018). Evidence from microstate studies suggests that altered resting-state brain network dynamics may represent a marker of risk to develop neuropsychiatric disorders (Tomescu et al., 2014, 2015; Andreou et al., 2014), predict clinical variables of an illness (Gschwind et al., 2016), or help to assess the efficacy of a treatment (Atluri et al., 2018; Sverak et al., 2018). Only two studies investigated resting-state EEG in BD patients (Strik et al., 1995; Damborská et al., 2019). These studies examined patients during a depressive episode within different affective disorders. Adaptive segmentation of resting-state EEG showed abnormal microstate topographies and reduced overall average microstate duration in patients that met criteria for unipolar or bipolar mood disorders or for dysthymia (Strik et al., 1995). Using a *k*-means cluster analysis, an increased occurrence of microstate A with depression as an effect related to the symptom severity was observed during a period of depression in unipolar and bipolar patients (Damborská et al., 2019).

Trait markers of BD based on neurobiological findings can be considered as biomarkers of illness (Piguet et al., 2016; Berchio et al., 2017). These trait markers of BD can be studied during the periods of remission, or euthymia. No microstate study on spontaneous activity, however, has been performed on euthymic BD patients to the best of our knowledge. Thus, the main goal of the current study was to explore group differences between euthymic patients with BD and healthy controls in terms of resting-state EEG microstate dynamics. We hypothesized that BD patients during remission will show altered temporal characteristics of EEG microstates such as duration, coverage, and occurrence.

## 2 Materials and Methods

### 2.1 Subjects

Data were collected from 17 euthymic adult patients with BD and 17 healthy control (HC) subjects. The patients were recruited from the Mood Disorders Unit at the Geneva University Hospital. A snowball convenience sampling was used for the selection of the BD patients. Control subjects were recruited by general advertisement. All subjects were clinically evaluated using clinical structured interview (DIGS: Diagnostic for Genetic Studies, (Nurnberger et al., 1994). BD was confirmed in the experimental group by the usual assessment of the specialized program, an interview with a psychiatrist, and a semi-structured interview and relevant questionnaires with a psychologist. Exclusion criteria for all participants were a history of head injury, current alcohol or drug abuse. Additionally, a history of psychiatric or neurological illness and of any neurological comorbidity were exclusion criteria for controls and bipolar patients, respectively. Symptoms of mania and depression were evaluated using the Young Mania Rating Scale (YMRS) (Young et al., 1978) and the Montgomery-Åsberg Depression Rating Scale (MADRS) (Williams and Kobak, 2008), respectively. Participants were considered euthymic if they scored < 6 on YMRS and < 12 on MADRS at the time of the experiment, and were stable for at least 4 weeks before. All patients were medicated, receiving pharmacological therapy including antipsychotics, antidepressants and mood stabilizers, and had to be under stable medication for at least 4 weeks. The experimental group included both BD I (*n* = 10) and BD II (*n* = 7) types.

To check for possible demographic or clinical differences between groups, subject characteristics such as age, education or level of depression were compared between groups using independent *t*-tests. Anxiety is highly associated with BD (Simon et al., 2004; 2007) and is a potential confounding variable when investigating microstate dynamics at rest. For example, decreased duration of EEG microstates at rest in patients with panic disorder has been reported (Wiedemann et al., 1998). To check for possible differences in anxiety symptoms, all subjects were assessed with the State-Trait Anxiety Inventory (STAI) (Spielberger et al., 1970). Anxiety as an emotional state (state-anxiety) and anxiety as a personal characteristic (trait-anxiety) were evaluated separately. Scores of both state- and trait-anxiety range from 20 to 80, higher values indicating greater anxiety. The scores were compared between patients and controls using independent *t*-tests.

This study was carried out in accordance with the recommendations of the Ethics Committee for Human Research of the Geneva University Hospital, with written informed consent from all subjects. All subjects gave written informed consent in accordance with the Declaration of Helsinki. The protocol was approved by the Ethics Committee for Human Research of the Geneva University Hospital, Switzerland.

### 2.2 EEG recording and pre-processing

The EEG was recorded with a high density 256-channel system (EGI System 200; Electrical Geodesic Inc., OR, USA), sampling rate of 1kHz, and Cz as acquisition reference. Subjects were sitting in a comfortable upright position and were instructed to stay as calm as possible, to keep their eyes closed and to relax for 6 minutes. They were asked to stay awake.

To remove muscular artifacts originating in the neck and face the data were reduced to 204 channels. Two to four minutes of EEG data were selected based on visual assessment of the artifacts and band-pass filtered between 1 and 40 Hz. Subsequently, in order to remove ballistocardiogram and oculo-motor artifacts, infomax-based Independent Component Analysis (Jung et al., 2000) was applied on all but one or two channels rejected due to abundant artifacts. Only components related to physiological noise, such as ballistocardiogram, saccadic eye movements, and eye blinking, were removed based on the waveform, topography and time course of the component. The cleaned EEG recordings were down-sampled to 125 Hz and the previously identified noisy channels were interpolated using a three-dimensional spherical spline (Perrin et al., 1989), and re-referenced to the average reference. All the preprocessing steps were done using MATLAB and the freely available Cartool Software 3.70 (https://sites.google.com/site/cartoolcommunity/home), programmed by Denis Brunet.

### 2.3 EEG data analysis

To estimate the optimal set of topographies explaining the EEG signal, a standard microstate analysis was performed using *k*-means clustering (see Supplementary Fig. 1). The polarity of the maps was ignored in this procedure (Brunet et al., 2011; Murray et al, 2008; Pascual-Marqui et al., 1995). To determine the optimal number of clusters, we applied a meta-criterion that is a combination of seven independent optimization criteria (for details see Bréchet et al., 2019). In order to improve the signal-to-noise ratio, only the data at the time points of the local maximum of the Global Field Power (GFP) were clustered (Pascual-Marqui et al., 1995; Koenig et al., 2002; Britz et al, 2010, Tomescu et al., 2014). The GFP is a scalar measure of the strength of the scalp potential field and is calculated as the standard deviation of all electrodes at a given time point (Michel et al., 1993; Brunet et al., 2011; Murray et al., 2008). The cluster analysis was first computed at the individual level yielding one set of representative maps for each subject. Clustering at the group level followed, in which the individual representative maps of patients and controls were clustered separately. Then all participants’ representative maps were clustered at global level yielding one set of maps that represented the data of all subjects. This one set of global representative maps entered the fitting process and the presence of each global map in every subject was determined. This enabled us to compare groups in terms of the presence of these global maps in the original data, i. e. to compare the microstate temporal characteristics between patients and controls.

In order to retrieve the temporal characteristics of the microstates, the fitting procedure consisted in calculating the spatial correlation between every representative map identified at the global level across all subjects and the individual subject’s scalp potential map in every instant of the pre-processed EEG recording. Each continuous time point of the subject’s EEG (not only the GFP peaks) was then assigned to the microstate class of the highest correlation (winner-takes-all), again ignoring polarity (Brunet et al., 2011; Bréchet et al., 2019; Michel and Koenig, 2018; Santarnecchi et al., 2017). Temporal smoothing parameters (window half size = 3, strength (Besag Factor) = 10) ensured that the noise during low GFP did not artificially interrupt the temporal segments of stable topography (Brunet et al., 2011; Pascual-Marqui et al., 1995). These segments of stable topography assigned to a given microstate class were then evaluated. For each subject, three temporal parameters were calculated for each microstate class: (i) occurrence, (ii) coverage, and (iii) duration. Occurrence indicates how many temporal segments of a given microstate class occur in one second. The coverage in percent represents the summed amount of time spent in a given microstate class as a portion of the whole analyzed period. The duration in milliseconds indicates the most common amount of time that a given microstate class is continuously present. The global explained variance for a specific microstate class was calculated by summing the squared spatial correlations between the representative map and its corresponding assigned scalp potential maps at each time point weighted by the GFP (Murray et al., 2008). Global explained variances of all microstate classes were then summed yielding the portion of data explained with the set of representative maps.

Microstate analysis was performed using the freely available Cartool Software 3.70, (https://sites.google.com/site/cartoolcommunity/home), programmed by Denis Brunet. Mann-Whitney U test was used to investigate group differences for temporal parameters of each microstate. Multiple comparisons were corrected using the false discovery rate (FDR) method (Benjamini, 2010).

Spearman’s rank correlations were calculated between the MADRS, YMRS, STAI-state, and STAI-trait scores and significant microstate parameters to check for possible relationships between symptoms and microstate dynamics. Eyes closed spontaneous EEG dynamics are mostly dominated by alpha activity (Niedermeyer, 2011). To provide an estimate of the absolute alpha power of each group, individual EEG data were Fourier transformed (Hanning windowing function, common averge reference). Mann-Whitney U test was used to investigate group differences for mean alpha (8-14Hz) power across 204 channels. To evaluate the role of the alpha rhythm in the appearance of the microstates, the Spearman’s rank correlation was calculated between Alpha Power and those microstate parameters, in which significant group differences were found. All statistical evaluations were performed by the routines included in the program package Statistica’13 (1984-2018, TIBCO, Software Inc, Version 13.4.0.14).

## 3 Results

### 3.1 Clinical and demographic variables

There were no significant differences in age and level of education between the patient and the control groups. In both groups, very low mean scores on depression and mania symptoms were observed, which did not significantly differ between the two groups. BD patients showed higher scores on state and trait scales of the STAI. For all subject characteristics, see Table 1.

**Table 1.**
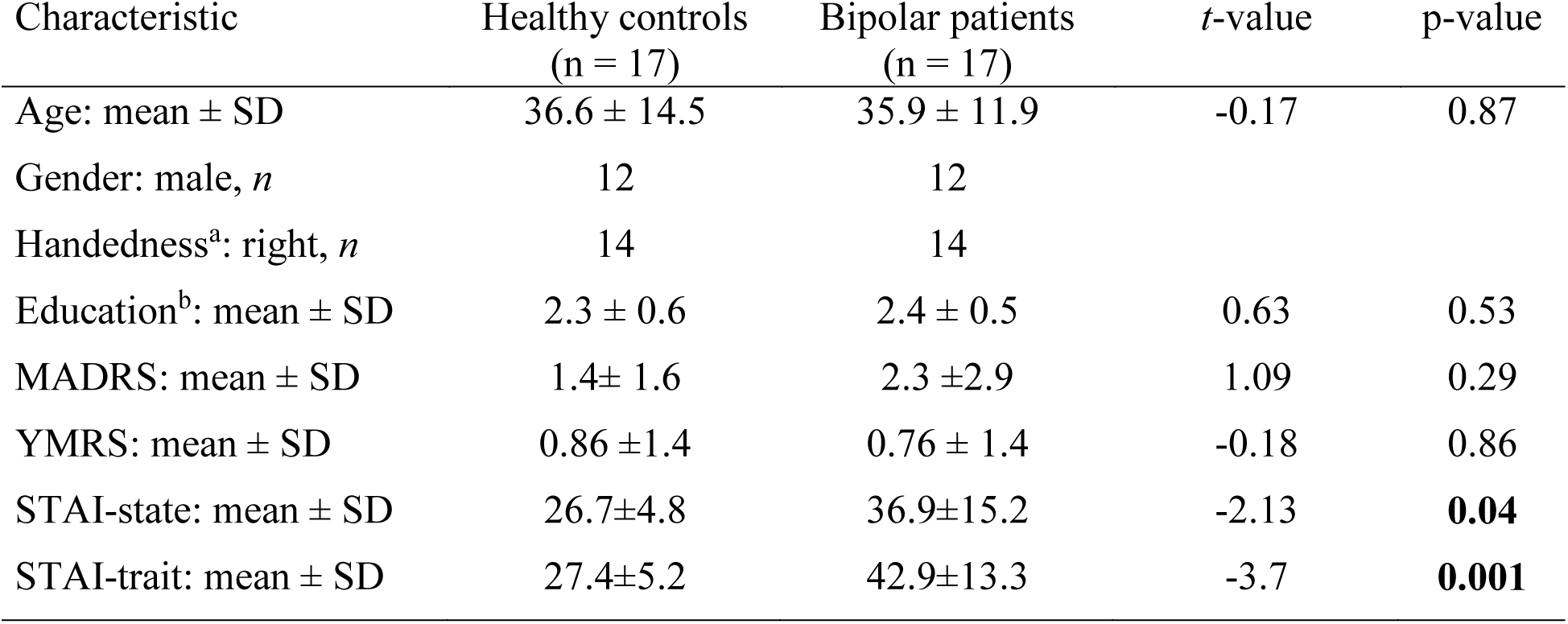
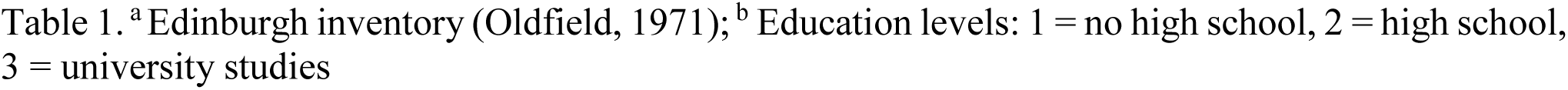
Subject characteristics

| Characteristic | Healthy controls<br>(n = 17) | Bipolar patients<br>(n = 17) | *t*-value | p-value |
| --- | --- | --- | --- | --- |
| Age: mean ± SD | 36.6 ± 14.5 | 35.9 ± 11.9 | -0.17 | 0.87 |
| Gender: male, *n* | 12 | 12 |  |  |
| Handedness<sup>a</sup>: right, *n* | 14 | 14 |  |  |
| Education<sup>b</sup>: mean ± SD | 2.3 ± 0.6 | 2.4 ± 0.5 | 0.63 | 0.53 |
| MADRS: mean ± SD | 1.4± 1.6 | 2.3 ±2.9 | 1.09 | 0.29 |
| YMRS: mean ± SD | 0.86 ±1.4 | 0.76 ± 1.4 | -0.18 | 0.86 |
| STAI-state: mean ± SD | 26.7±4.8 | 36.9±15.2 | -2.13 | **0.04** |
| STAI-trait: mean ± SD | 27.4±5.2 | 42.9±13.3 | -3.7 | **0.001** |

### 3.2 Microstate results

The meta-criterion used to determine the most dominant topographies revealed five resting-state microstate maps across patients, healthy controls, and all subjects, explaining 82.1 %, 83.1 %, and 82.2 % of the global variance, respectively (Fig. 1). The topographies resembled those previously reported as A, B, C, and D maps (Khanna et al., 2015; Michel and Koenig, 2018; Koenig et al., 2002; Britz et al., 2010) and one of the three recently identified additional maps (Custo et al., 2017). We labeled these scalp maps from A to E in accordance with the previous literature on microstates. The scalp topographies showed left posterior-right anterior orientation (map A), a right posterior-left anterior orientation (map B), an anterior-posterior orientation (map C), a fronto-central maximum (map D), and a parieto-occipital maximum (map E).

**Figure 1.**
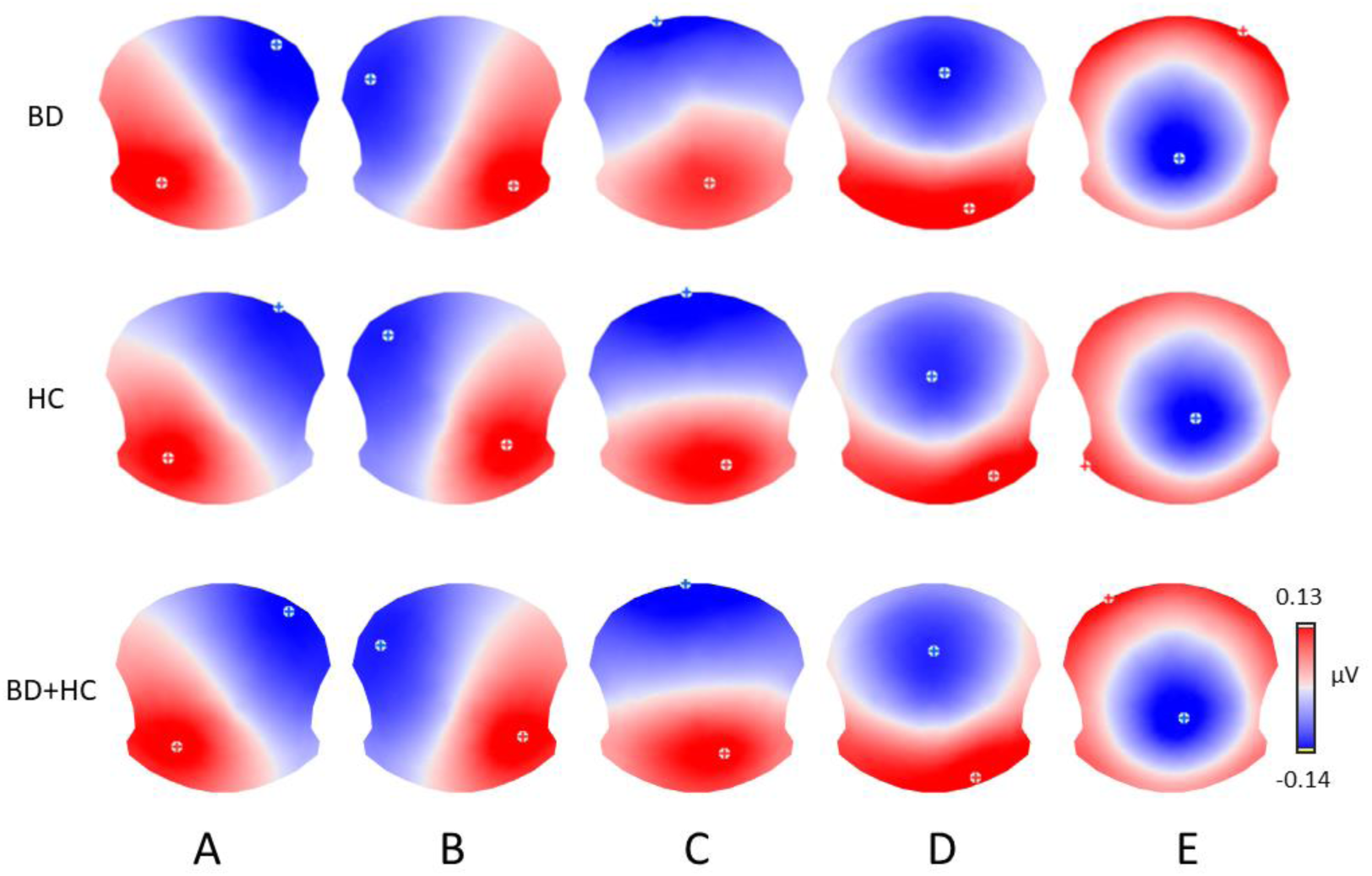
The five microstate topographies identified in the group clusterings across patients (BD) and healthy controls (HC) and in the global clustering across all subjects (BD+HC). Note that similar five dominant microstate topographies were identified in all three clusterings.

Since some microstate parameters showed a non-homogeneity of variances in the two groups (Levene’s tests for the microstate C coverage and microstates A and C duration; p<0.01), we decided to calculate Mann-Whitney U test to investigate group differences for temporal parameters of each microstate.

We found significant between-group differences for microstate classes A and B. Both microstates showed increased presence in patients in terms of occurrence and coverage. The two groups did not differ in any temporal parameter of microstates C, D, or E. The results of the temporal characteristics of each microstate are summarized in Table 2 and Fig. 2.

**Table 2.**
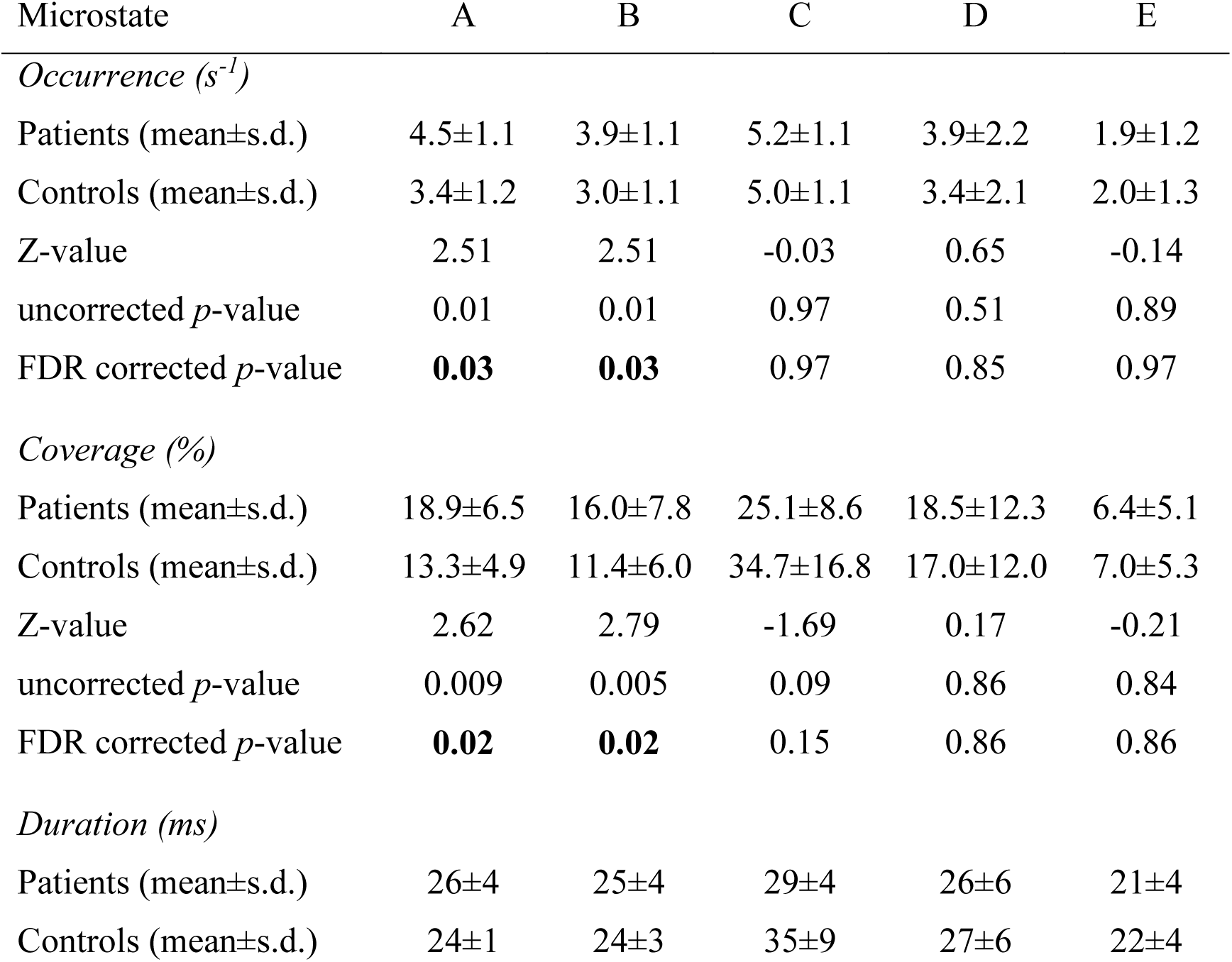
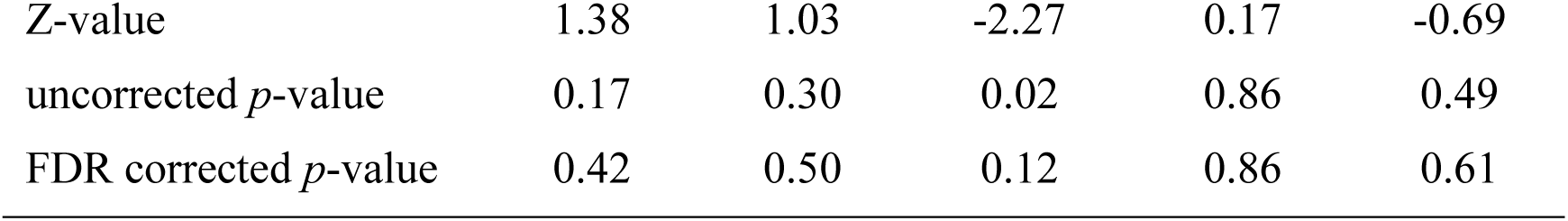
Mann-Whitney U test for group comparisons in the investigated microstate parameters

| Microstate | A | B | C | D | E |
| --- | --- | --- | --- | --- | --- |
| *Occurrence (s<sup>-1</sup>)* |  |  |  |  |  |
| Patients (mean±s.d.) | 4.5±1.1 | 3.9±1.1 | 5.2±1.1 | 3.9±2.2 | 1.9±1.2 |
| Controls (mean±s.d.) | 3.4±1.2 | 3.0±1.1 | 5.0±1.1 | 3.4±2.1 | 2.0±1.3 |
| Z-value | 2.51 | 2.51 | -0.03 | 0.65 | -0.14 |
| uncorrected *p*-value | 0.01 | 0.01 | 0.97 | 0.51 | 0.89 |
| FDR corrected *p*-value | **0.03** | **0.03** | 0.97 | 0.85 | 0.97 |
| *Coverage (%)* |  |  |  |  |  |
| Patients (mean±s.d.) | 18.9±6.5 | 16.0±7.8 | 25.1±8.6 | 18.5±12.3 | 6.4±5.1 |
| Controls (mean±s.d.) | 13.3±4.9 | 11.4±6.0 | 34.7±16.8 | 17.0±12.0 | 7.0±5.3 |
| Z-value | 2.62 | 2.79 | -1.69 | 0.17 | -0.21 |
| uncorrected *p*-value | 0.009 | 0.005 | 0.09 | 0.86 | 0.84 |
| FDR corrected *p*-value | **0.02** | **0.02** | 0.15 | 0.86 | 0.86 |
| *Duration (ms)* |  |  |  |  |  |
| Patients (mean±s.d.) | 26±4 | 25±4 | 29±4 | 26±6 | 21±4 |
| Controls (mean±s.d.) | 24±1 | 24±3 | 35±9 | 27±6 | 22±4 |
| Z-value | 1.38 | 1.03 | -2.27 | 0.17 | -0.69 |
| uncorrected *p*-value | 0.17 | 0.30 | 0.02 | 0.86 | 0.49 |
| FDR corrected *p*-value | 0.42 | 0.50 | 0.12 | 0.86 | 0.61 |

**Figure 2.**
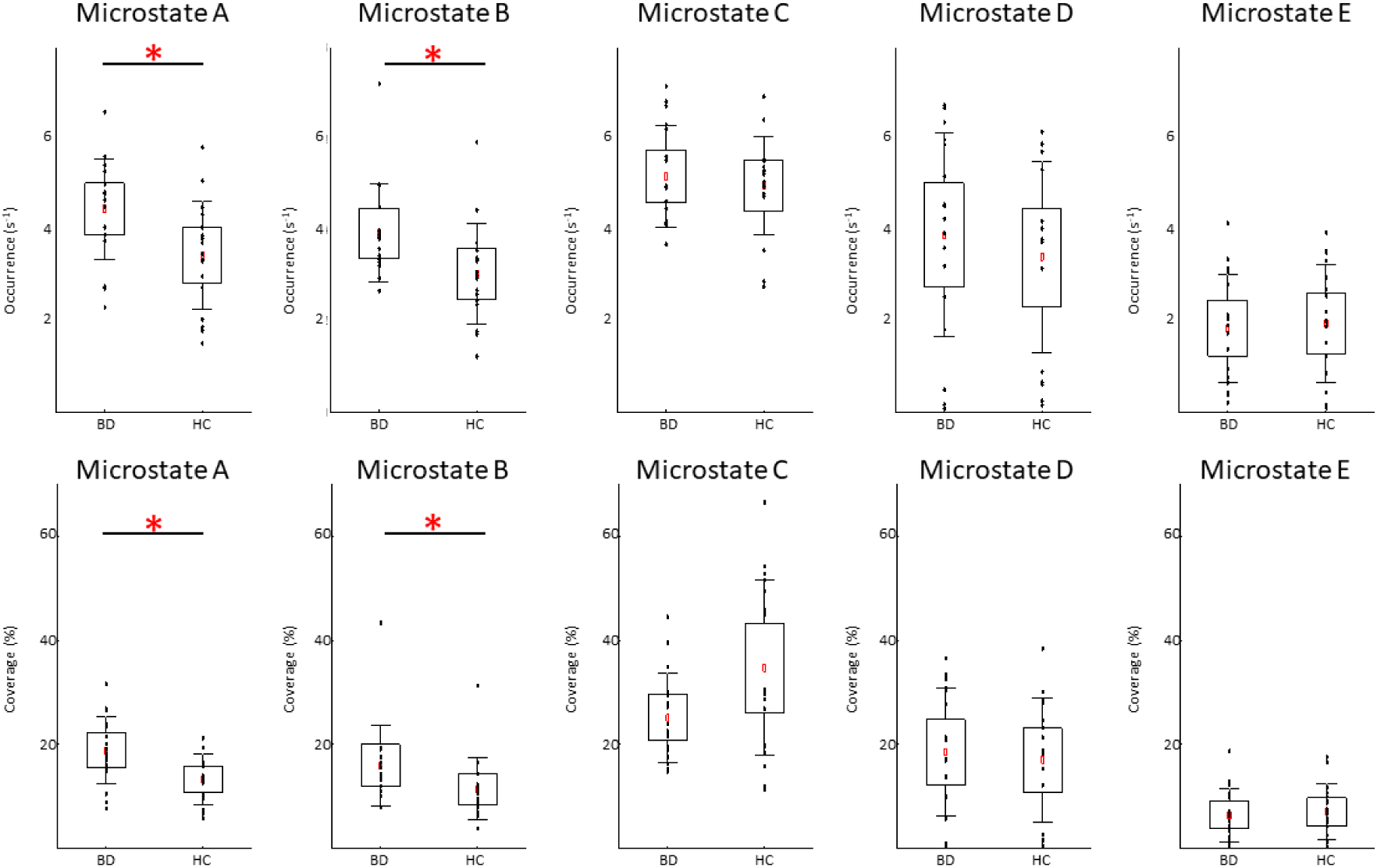
Temporal dynamics of EEG microstates in patients with bipolar disorder (BD) and in healthy controls (HC). In each subplot, the raw data are plotted on top of the boxplot showing the mean 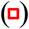, 95% confidence interval (box plot area), 1 standard deviation (whiskers), and significant differences (*). In all plots, x-axes represent the subject group; y-axes represent the occurrence (upper plots) or coverage (lower plots). Note significantly increased occurrence and coverage of the microstate A and B in the BD compared to HC group (FDR corrected p < 0.05).

### 3.3 Clinical correlations

Correlations with clinical parameters were calculated for those microstate parameters, in which significant group differences were found. The results of Spearman’s rank correlation revealed a significant positive association between the coverage of the microstate B and the STAI-state (r = 0.40; p<0.05) and STAI-trait (r = 0.54; p<0.05) scores. The results of Spearman’s rank correlation revealed a significant positive association between the occurrence of the microstate B and the STAI-trait (r = 0.47; p<0.05) scores. The results of Spearman’s rank correlation revealed no significant associations between the STAI-state or STAI-trait scores and the occurrence or coverage of the microstate A (all absolute r-values < 0.35).

The results of Spearman’s rank correlation revealed no significant associations between the MADRS and YMRS scores and the occurrence or coverage of the microstate A and B (all absolute r-values < 0.30).

### 3.4 Alpha rhythm

The Mann-Whitney U test showed significantly decreased alpha power (p < 0.03, Z-value 2.7) in the BD compared to HC group (see Fig. 3). The results of Spearman’s rank correlation revealed no significant associations between the alpha power and occurrence or coverage of microstates A and B (all absolute r-values < 0.40).

**Figure 3.**
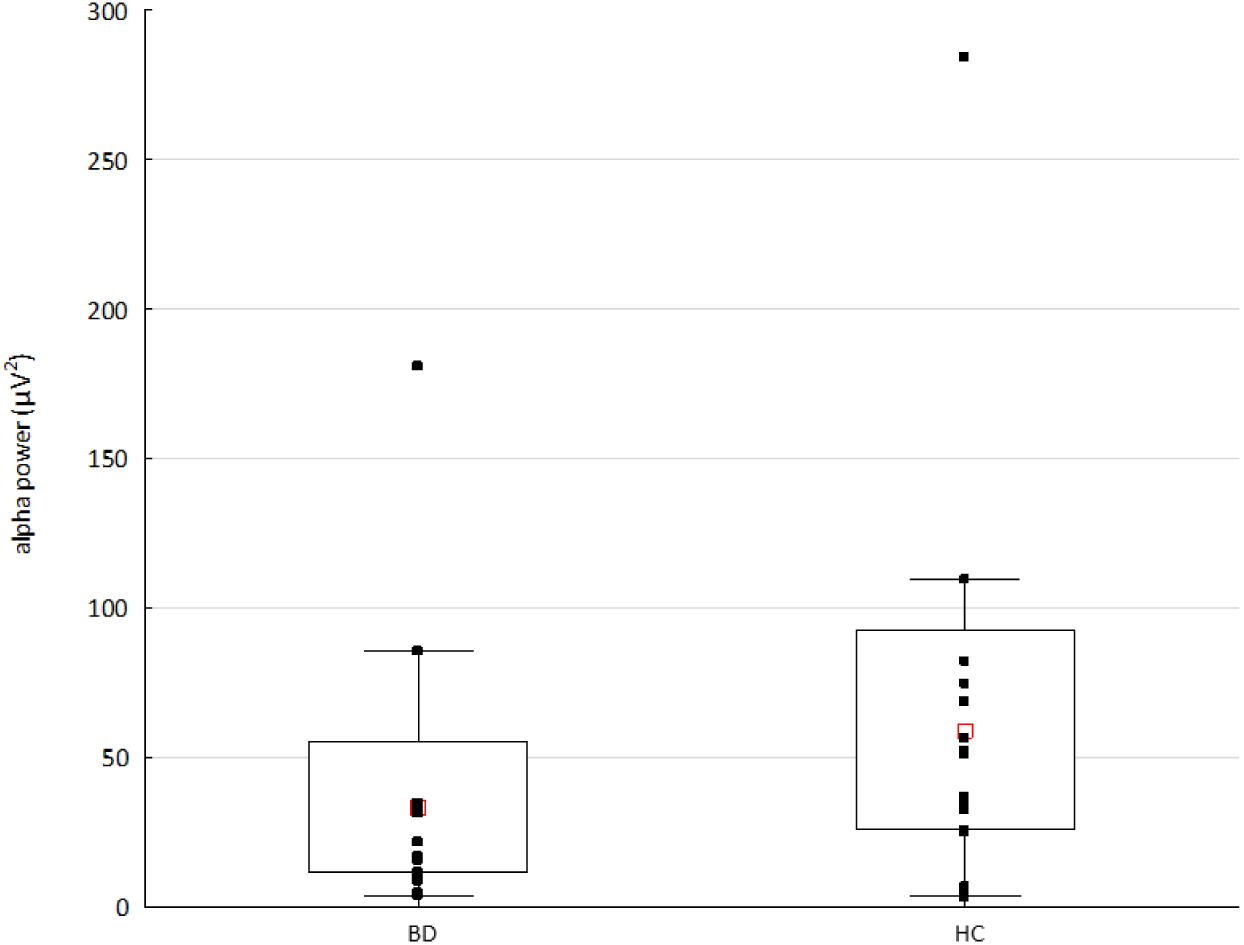
The alpha power in patients with bipolar disorder (BD) and healthy controls (HC). Raw data are plotted on top of each boxplot showing the mean 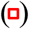, 95% confidence interval (box plot area), and non-outlier range (whiskers). The x-axis represents the subject group; the y-axis represents the average alpha (8-14 Hz) power across 204 channels. Note significantly decreased alpha power in the BD compared to HC group (p < 0.03, Z value 2.7).

## 4 Discussion

Our study presents the first evidence for altered resting-state EEG microstate dynamics in euthymic patients with BD. Patients were stable and did not significantly differ in their depressive or manic symptomatology from healthy controls at the time of experiment. Despite this fact, they showed abnormally increased presence of microstates A and B, the latter correlating with the anxiety level.

In an earlier combined fMRI-EEG study the microstate A was associated with the auditory network (Britz et al., 2010). Moreover, generators of the functional EEG microstates were estimated in recent studies, where sources of the microstate A showed left-lateralized activity in the temporal lobe, insula, mPFC, and occipital gyri (Custo et al., 2017; Bréchet et al., 2019).

In the fMRI literature as well, resting-state functional connectivity alterations of the insula (Yin et al., 2018), the auditory network (Reinke et al., 2013), and the mPFC (Gong et al., 2019) were reported in BD patients. Verbal episodic memory deficits and language-related symptoms in BD patients were suggested to be associated with a diminished functional connectivity within the auditory/temporal gyrus and to be compensated by increased fronto-temporal functional connectivity (Reinke et al., 2013). The mPFC was also identified as a major locus of shared abnormality in BD and schizophrenia (Öngür et al., 2010), showing reduced default mode network connectivity from the mPFC to the hippocampus and fusiform gyrus, as well as increased connectivity between the mPFC and primary visual cortex in BD. Hypoconnectivity of the default mode network from the left posterior cingulate cortex to the bilateral mPFC and bilateral precuneus, and reduced salience connectivity of the left sgACC to the right inferior temporal gyrus in BD patients (Gong et al., 2019) was observed in unmedicated BD patients. In euthymic BD subjects compared to healthy controls, resting-state functional connectivity of the insula (Minuzzi et al., 2018) and amygdala (Li et al., 2018) to other brain regions was reported to be increased and decreased, respectively. In summary, the evidence from fMRI studies shows both hypoconnectivity (Gong et al., 2019; Öngür et al., 2010) and hyperconnectivity (Minuzzi et al., 2018; Reinke et al., 2013; Öngür et al., 2010) pointing to complex alterations of functional resting-state networks. Our findings of increased presence of the microstate A in euthymic BD patients might be related to the hyperconnectivity of the underlying networks that involve the temporal lobe, insula, mPFC, and occipital gyri.

Anxiety symptoms were previously associated with greater severity and impairment in BD (Simon et al., 2004) and euthymic bipolar patients tend to present high residual level of anxiety (Albert et al., 2008), as it was the case here. No significant correlation was found between the increased anxiety scores and the increased occurrence or coverage of the microstate A. Our results therefore indicate that this alteration of microstate dynamics might represent a characteristic feature of BD that is not affected by anxiety. The demonstrated alterations in microstate A dynamics during clinical remission might reflect (i) an impaired resting-state large-scale brain network dynamics as a trait characteristic of the disorder and/or (ii) a compensatory mechanism needed for clinical stabilization of the disorder.

Our study is the first to examine EEG microstate dynamics of spontaneous activity in BD patients during remission. The here demonstrated group difference of BD patients vs. controls, is not congruent with the previously reported reduced duration of the EEG microstates during a depressive episode (Strik et al., 1995). The experimental group in that study was not restricted to bipolar patients, however, and included also patients who met the criteria for unipolar depression or dysthymia. Moreover, authors examined the overall microstate duration and did not examine distinct microstates separately. These and other aspects, such as different clustering methods used, make it difficult to compare our findings with that early evidence of disrupted microstate dynamics in depression. Nevertheless, in our recent resting-state study we showed positive associations of depressive symptoms with the occurrence of microstate A in a heterogenous group of patients with affective disorders (Damborská et al., 2019).

The microstate B was previously associated with the visual network (Britz et al., 2010; Custo et al., 2014, 2017; Bréchet et al., 2019) and thoughts related to the conscious experience of an episodic autobiographic memory, i.e. mental visualization of the scene (Bréchet et al., 2019). In our group of BD patients, we found an abnormally increased presence of microstate B that was associated with a higher anxiety. In particular, the occurence together with coverage and only the coverage were positively correlated with scores of trait and state anxiety, respectively. The observed change in microstate B dynamics might be, therefore, more related to a relatively stable disposition than to the actual emotional state. Previous studies also suggest that anxiety may influence visual processing (Phelps et al., 2006; Laretzaki et al., 2010) and that connections between amygdala and visual cortex might underlie enhanced visual processing of emotionally salient stimuli in patients with social fobia (Goldin et al., 2009). Our finding of increased presence of microstate B positively associated with anxiety level in euthymic BD patients is consistent with these observations. Additionally, a more regular appearance for microstate B and increased overall temporal dependencies among microstates were recently reported in mood and anxiety disorders, suggesting a decreased dynamicity in switching between different brain states in these psychiatric conditions (Al Zoubi et al., 2019). Another microstate study on anxiety disorders reported a decreased overall resting-state microstate duration in panic disorder (Wiedemann et al., 1998). That early study, however, did not assess temporal characteristics of different microstates separately and it is therefore difficult to compare those findings with our observations. Further evidence is needed to determine, whether the increased presence of microstate B in our experimental group is a characteristic feature of BD or anxiety, or whether it is related to both conditions.

We found an unchanged duration but a higher occurence and coverage of A and B microstates in BD patients. In other words, an unchanged sustainability in time and still increased presence of these microstates in patients compared to healthy controls were observed. Possible explanation for this finding appears to be a redundance in activation of the sensory and autobiographic memory networks during spontaneous mentation in euthymic BD patients. Changes in A and B microstate occurence, duration, and coverage have been reported in several psychiatric conditions such as schizophrenia, dementia, narcolepsy, multiple sclerosis, panic disorder, etc. (for review see Michel and Koenig, 2018). In schizophrenia patients, both increased (Andreou et al., 2014) and decreased (Nishida et al., 2013; Lehmann et al., 2005) durations of microstate B as well as increased occurrence of microstate A (Lehmann et al., 2005) were reported. Increases in duration and occurrence of microstates A and B were also observed in patients with multiple sclerosis, moreover predicting depression scores and other clinical variables (Gschwind, et al., 2016). It was suggested that multiple sclerosis affects the “sensory” (visual, auditory) rather than the higher-order (salience, central executive) functional networks (Michel and Koenig, 2018). Our findings of impaired dynamics in microstates A and B suggest a similar interpretation for the BD. Although the pathophysiology in multiple sclerosis differes from that in depression, the increased presence of A and B microstates might reflect aberrant temporal functioning of sensory-related and memory networks in both diseases. Evidence from fMRI studies points to topographical dysbalances between the default mode and sensorimotor networks in BD patients with opposing patterns in depression and mania (Martino et al., 2016). Cyclothymic and depressive temperaments were associated with opposite changes in the sensorimotor network variability in the resting state signal measured by fractional standard deviation of Blood-Oxygen-Level Dependent signal (Conio et al., 2019). Our findings of altered microstates A and B dynamics is consistent with this fMRI evidence of impaired sensorimotor network in affective disorders, and moreover suggests that neural correlates of these deficits are prominent even during the euthymic state in BD patients.

It is known that EEG microstate dynamics during eyes closed and eyes open resting-state is different. Manipulations of visual input showed increased occurrence and coverage of microstate B and shorter duration of microstate A in the eyes open condition in healthy young adults (Seitzman et al. 2017). Further studies are needed to investigate, whether the here observed abnormalities of the A and B microstates in BD patients are still present in the eyes open condition.

BD patients were previously shown to display lower alpha power as compared to healthy controls (Basar et al., 2012), as it was the case here. We failed, however, to find any significant correlation between the altered microstate dynamics and decreased alpha power. Our findings, therefore, further support the previously reported independence of microstates from EEG frequency power fluctuations (Britz et al., 2010).

In summary, results of the current study seem to indicate that dysfunctional activity of resting-state brain networks underlying microstates A and B is a detectable impairment in BD during an euthymic state. The presence of microstate A and B represents measures that might be implicated in clinical practice, although using these parameters for early identification of BD at individual level could prove challenging. If future studies confirm the same pattern in prodromal or vulnerable subjects, it could help detection of at-risk subjects and therefore the possiblility for early intervention. The present study has, however, some limitations. Our low sample size made it impossible to examine any potential influence of medication on the microstate parameters by comparing patients receiving a specific drug with those not receiving it. Possible effects of medication on our results should be therefore taken into account. Due to the same reason, it was not possible to examine any potential influence of subtypes of BD on microstate results.

## 5 Conclusions

Our study described altered EEG resting-state microstate temporal parameters in euthymic bipolar patients. Our findings provide an insight into the resting-state global brain network dynamics in BD. Since the increased presence of microstate A is not unique to BD patients, having been reported also in other psychiatric disorders (see Michel and Koenig, 2018), it might be considered only as a non-specific electrophysiological marker of BD. Moreover, studies examining possible interactions between microstate dynamics and BD symptoms are needed to better understand the dysfunction of large-scale brain network resting-state dynamics in this affective disorder.

---

**Supplementary Figure 1.**
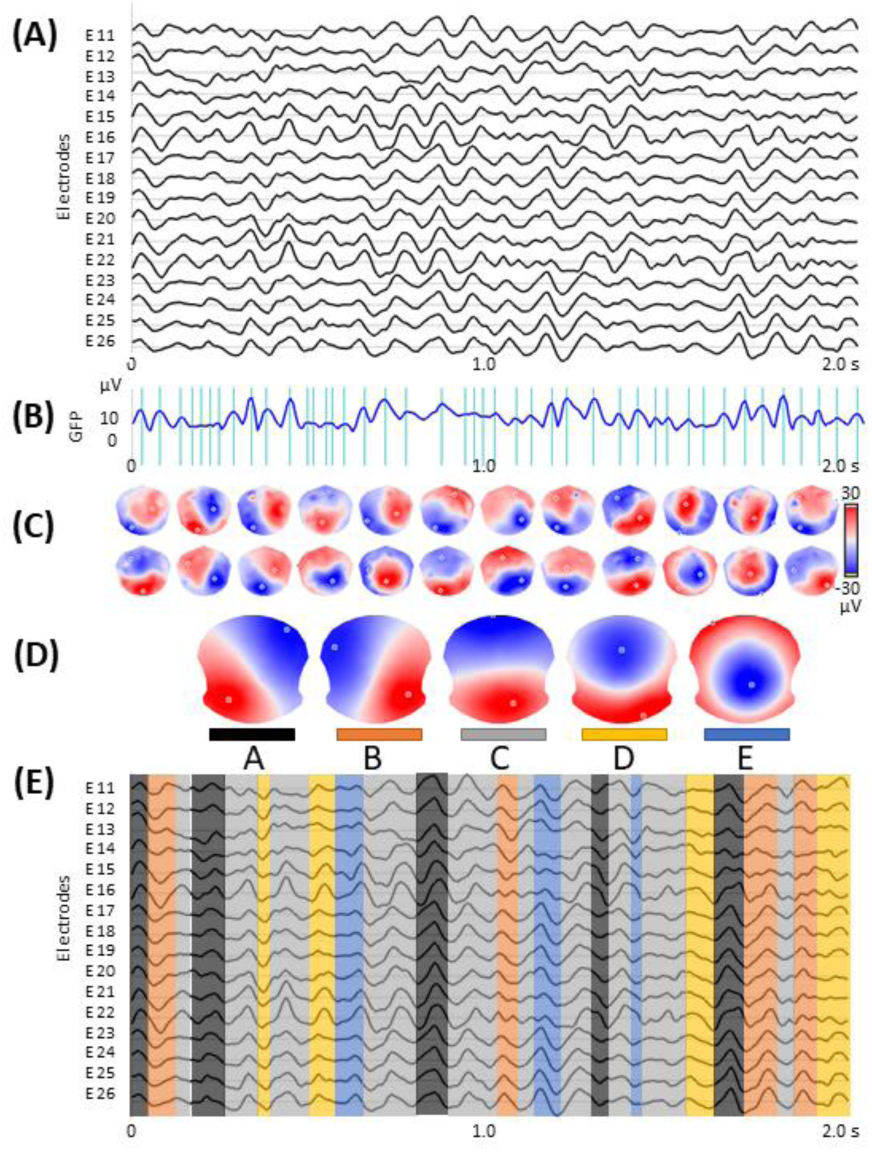
Microstate analysis: **(A)** resting-state EEG from subsample of 16 out of 204 electrodes; **(B)** global field power (GFP) curve with the GFP peaks (vertical lines) in the same EEG period as shown in (A); **(C)** potential maps at successive GFP peaks, indicated in (B), from the first 1 s period of the recording; **(D)** set of five cluster maps best explaining the data as revealed by K-means clustering of the maps at the GFP peaks; **(E)** the original EEG recording shown in (A) with superimposed color-coded microstate segments. Note that each time point of the EEG recording was labelled with the cluster map, shown in (D), with which the instant map correlated best. The duration of segments, occurrence, and coverage for all microstates were computed on thus labeled EEG recording.

## 10 Author Contributions

AD – designed the study, performed the analysis, and wrote the initial draft; JMA, AGD and CP – were responsible for clinical assessment; CMM – served as an advisor; CB – collected the HD-EEG data and was responsible for the overall oversight of the study. All authors revised the manuscript.

## 12 Acknowledgments

Special thanks go to Anne-Lise Kung, psychologist, for her involvement in clinical data collection.

## 14 Data Availability Statement

The raw data supporting the conclusions of this manuscript will be made available by the authors, without undue reservation, to any qualified researcher.

## References

Albert, U., Rosso, G., Maina, G., Bogetto, F., (2008). Impact of anxiety disorder comorbidity on quality of life in euthymic bipolar disorder patients: differences between bipolar I and II subtypes. J. Affective Disord. 105, 297–303.

Al Zoubi, O., Mayeli, A., Tsuchiyagaito, A., Misaki, M., Zotev, V., Refai, H., et al. (2019). EEG microstates temporal dynamics differentiate individuals with mood and anxiety disorders from healthy subjects. Front. Human Neurosci.;13.

Andreou, C., Faber, P.L., Leicht, G., Schoettle, D., Polomac, N., Hanganu-Opatz, I.L., et al. (2014). Resting-state connectivity in the prodromal phase of schizophrenia: Insights from EEG microstates. Schizophr. Res. 152, 513–520.

Atluri, S., Wong, W., Moreno, S., Blumberger, D.M., Daskalakis, Z.J., Farzan, F., (2018). Selective modulation of brain network dynamics by seizure therapy in treatment-resistant depression. NeuroImage Clin. 20, 1176–1190.

Basar, E., Güntekin, B., Atagün, I., Turp Gölbas, B., Tülay, E., Özerdem, A. (2012). Brain’s alpha activity is highly reduced in euthymic bipolar disorder patients. Cogn. Neurodynamics 6, 11–20.

Benjamini Y., (2010). Discovering the false discovery rate. J. R. Stat. Soc. Ser. B Stat. Methodol. 72, 405–416.

Berchio C., Piguet C., Michel C.M., Cordera P., Rihs T.A., Dayer A.G., et al. (2017). Dysfunctional gaze processing in bipolar disorder. NeuroImage Clin. 16, 545–556.

Bréchet, L., Brunet, D., Birot, G., Gruetter, R., Michel C.M., Jorge, J., (2019). Capturing the spatiotemporal dynamics of self-generated, task-initiated thoughts with EEG and fMRI. Neuroimage, 194, 82–92.

Bressler, S.L., Menon, V., (2010) Large-scale brain networks in cognition: emerging methods and principles. Trends Cogn. Sci. 14, 277–290.

Britz, J., Van De Ville D., Michel, C.M., (2010). BOLD correlates of EEG topography reveal rapid resting-state network dynamics. NeuroImage, 52, 1162–1170.

Brunet, D., Murray, M.M., Michel, C.M., (2011). Spatiotemporal analysis of multichannel EEG: CARTOOL. Comput. Intell. Neurosci. 813870

Changeux, J.P. and Michel, C.M., (2004). Mechanism of neural Integration at the Brain-scale Level. In: Grillner, S., Graybiel, A.M. (Eds.), Microcircuits. MIT Press, Cambridge, pp. 347–370.

Chase, H.W., Phillips, M.L. (2016) Elucidating Neural Network Functional Connectivity Abnormalities in Bipolar Disorder: Toward a Harmonized Methodological Approach. Biol. Psychiatry Cogn. Neurosci. Neuroimaging 1, 288–298.

Chen, C.H., Suckling, J., Lennox, B.R., Ooi, C., Bullmore, E.T., (2011). A quantitative meta-analysis of fMRI studies in bipolar disorder. Bipolar Disord. 13, 1–15.

Cloutier, M., Greene, M., Guerin, A., Touya, M., Wu, E., (2018). The economic burden of bipolar I disorder in the United States in 2015. J. Affective Disord., 226, 45–51.

Conio, B., Magioncalda, P., Martino, M., Tumati, S., Capobianco, L., Escelsior, A., et al. (2019). Opposing patterns of neuronal variability in the sensorimotor network mediate cyclothymic and depressive temperaments. Hum. Brain Mapp. 40, 1344–1352.

Custo, A., Vulliemoz, S., Grouiller, F., Van De Ville, D., Michel, C., (2014). EEG source imaging of brain states using spatiotemporal regression. Neuroimage 96, 106–116

Custo, A., Van De Ville, D., Wells, W.M., Tomescu, M.I., Brunet, D., Michel, C.M., (2017). Electroencephalographic Resting-State Networks: Source Localization of Microstates. Brain Connectivity 7, 671–682.

Damborská, A., Tomescu, M.I., Honzírková, E., Barteček, R., Hořínková, J., Fedorová, S. et al. (2019) EEG resting-state large-scale brain network dynamics are related to depressive symptoms. Frontiers in Psychiatry, 548 (10).

Eaton, W.W., Alexandre, P., Bienvenu, O.J., Clarke, D., Martins, S.S., Nestadt, G., et al. (2012). The Burden of Mental Disorders, in: Eaton W.W. (Ed.), Public Mental Health, Oxford University Press

Ferrari, A.J., Stockings, E., Khoo, J.P., Erskine, H.E., Degenhardt, L., Vos, T., et al. (2016). The prevalence and burden of bipolar disorder: findings from the Global Burden of Disease Study 2013. Bipolar Disord.; 18, 440–450.

Goldin, P.R., Manber, T., Hakimi, S., Canli, T., Gross, J.J. (2009). Neural bases of social anxiety disorder: Emotional reactivity and cognitive regulation during social and physical threat. Arch. Gen. Psychiatry 66, 170–180.

Gong J., Chen G., Jia Y., Zhong S., Zhao L., Luo X., et al. (2019). Disrupted functional connectivity within the default mode network and salience network in unmedicated bipolar II disorder. Prog. Neuro-Psychopharmacol. Biol. Psychiatry 88, 11–18.

Gschwind, M., Hardmeier, M., Van De Ville, D., Tomescu, M.I., Penner, I., Naegelin, Y., et al. (2016). Fluctuations of spontaneous EEG topographies predict disease state in relapsing-remitting multiple sclerosis. NeuroImage Clin. 12, 466–477.

Hirschfeld, R.M.A., (2007). Screening for bipolar disorder. Am. J. Managed Care 13, S164–S169.

Houenou, J., Frommberger, J., Carde, S., Glasbrenner, M., Diener, C., Leboyer, M., et al. (2011). Neuroimaging-based markers of bipolar disorder: Evidence from two meta-analyses. J. Affective Disord. 132, 344–355.

Jung, T., Makeig, S., Westerfield, M., Townsend, J., Courchesne, E.c Sejnowski, T. J., (2000). Removal of eye activity artifacts from visual event-related potentials in normal and clinical subjects. Clin. Neurophysiol. 111, 1745–1758.

Khanna, A., Pascual-Leone, A., Michel, C.M., Farzan F., (2015). Microstates in resting-state EEG: Current status and future directions. Neurosci. Biobehav. Rev. 49, 105–113.

Koenig, T., Prichep, L., Lehmann, D., Sosa, P.V., Braeker, E., Kleinlogel, H., et al. (2002). Millisecond by millisecond, year by year: Normative EEG microstates and developmental stages. Neuroimage 16, 41–48.

Kupferschmidt, D.A., Zakzanis, K.K., (2011). Toward a functional neuroanatomical signature of bipolar disorder: Quantitative evidence from the neuroimaging literature. Psychiatry Res. Neuroimaging 193, 71–79.

Laretzaki, G., Plainis, S., Argyropoulos, S., Pallikaris, I.G., Bitsios, P. (2010). Threat and anxiety affect visual contrast perception. J Psychopharmacol 24, 667–675.

Lehmann, D., Ozaki, H., Pal, I., (1987). EEG alpha map series: brain micro-states by space-oriented adaptive segmentation. Electroencephalogr. Clin. Neurophysiol. 67, 271–288.

Lehmann D., (1990). Brain Electric Microstates and Cognition: The Atoms of Thought. In: John E.R., Harmony T., Prichep L.S., Valdés-Sosa M., Valdés-Sosa P.A. (eds) Machinery of the Mind. Birkhäuser, Boston, MA, pp. 209–224.

Lehmann, D., Faber, P.L., Galderisi, S., Herrmann, W.M., Kinoshita, T., Koukkou, M., et al. (2005). EEG microstate duration and syntax in acute, medication-naïve, first-episode schizophrenia: A multi-center study. Psychiatry Res. Neuroimaging, 138, 141–156.

Li, G., Liu, P., Andari, E., Zhang, A., Zhang, K., (2018). The role of amygdala in patients with euthymic bipolar disorder during resting state. Front. Psychiatry 9(SEP).

Martino, M., Magioncalda, P., Huang, Z., Conio, B., Piaggio, N., Duncan, N.W., et al. (2016). Contrasting variability patterns in the default mode and sensorimotor networks balance in bipolar depression and mania. Proc. Natl. Acad. Sci. U S A 113, 4824–4829.

Merikangas, K.R., Jin, R., He, J. P., Kessler, R.C., Lee, S., Sampson, N.A., et al. (2011). Prevalence and correlates of bipolar spectrum disorder in the World Mental Health Survey Initiative. Arch. Gen. Psychiatry 68, 241–251.

Michel, C.M., Brandeis, D., Skrandies, W., Pascual, R., Strik, W.K., Dierks, T. et al. (1993). Global field power: a ‘time-honoured’ index for EEG/EP map analysis. Int. J. Psychophysiol. 15, 1–2.

Michel, C.M. and Koenig, T., (2018). EEG microstates as a tool for studying the temporal dynamics of whole-brain neuronal networks: A review. Neuroimage 180, 577–593.

Minuzzi, L., Syan, S.K., Smith, M., Hall, A., Hall, G.B.C., Frey, B.N., (2018). Structural and functional changes in the somatosensory cortex in euthymic females with bipolar disorder. Aust. New Zealand J. Psychiatry 52, 1075–1083.

Murray, M.M., Brunet, D., Michel, C.M., (2008). Topographic ERP analyses: A step-by-step tutorial review. Brain Topogr. 20, 249–264.

Niedermeyer, E. Niedermeyer’s electroencephalography: basic principles, clinical applications, and related fields. Lippincott Williams & Wilkins, 2011.

Nishida, K., Morishima, Y., Yoshimura, M., Isotani, T., Irisawa, S., Jann, K., et al. (2013). EEG microstates associated with salience and frontoparietal networks in frontotemporal dementia, schizophrenia and Alzheimer’s disease. Clin. Neurophysiol. 124, 1106–1114.

Nurnberger Jr., J.I., Blehar, M.C., Kaufmann, C.A., York-Cooler, C., Simpson, S.G., Harkavy-Friedman, J., et al. (1994). Diagnostic interview for genetic studies: Rationale, unique features, and training. Arch. Gen. Psychiatry 51, 849–859.

Oldfield, R.C., (1971). The assessment and analysis of handedness: The Edinburgh inventory. Neuropsychologia 9, 97–113.

Öngür, D., Lundy, M., Greenhouse, I., Shinn, A.K., Menon, V., Cohen, B.M., et al. (2010). Default mode network abnormalities in bipolar disorder and schizophrenia. Psychiatry Res Neuroimaging 183, 59–68.

Pascual-Marqui, R.D., Michel, C.M., Lehmann, D., (1995). Segmentation of Brain Electrical Activity into Microstates; Model Estimation and Validation. IEEE Trans. Biomed. Eng. 42, 658–665.

de Pasquale. F., Corbetta, M., Betti. V., Della Penna, S., (2018). Cortical cores in network dynamics. Neuroimage 180, 370–382.

Perrin, F., Pernier, J., Bertrand, O., Echallier, J. F., (1989). Spherical splines for scalp potential and current density mapping. Electroencephalogr. Clin. Neurophysiol. 72, 184–187.

Phelps, E.A., Ling, S., Carrasco, M. (2006) Emotion facilitates perception and potentiates the perceptual benefits of attention. Psychol. Sci. 17, 292–299.

Piguet, C., Dayer, A., Aubry, J.M., (2016). Biomarkers and vulnerability to bipolar disorders. Schweiz Arch. Neurol. Psychiatr. 167, 57–67.

Piguet, C., Fodoulian, L., Aubry, J.M., Vuilleumier, P., Houenou, J., (2015). Bipolar disorder: Functional neuroimaging markers in relatives. Neurosci. Biobehav. Rev. 57, 284–296.

Reinke, B., Van de Ven, V., Matura, S., Linden, D.E.J., Oertel-Knöchel, V. (2013). Altered intrinsic functional connectivity in language-related brain regions in association with verbal memory performance in euthymic bipolar patients. Brain Sci. 3, 1357–1373.

Rey, G., Piguet, C., Benders, A., Favre, S., Eickhoff, S.B., Aubry, J.M., et al. (2016). Resting-state functional connectivity of emotion regulation networks in euthymic and non-euthymic bipolar disorder patients. Eur. Psychiatry 34, 56–63.

Santarnecchi, E., Khanna, A.R., Musaeus, C.S., Benwell, C.S.Y., Davila, P., Farzan, F., et al. (2017). EEG Microstate Correlates of Fluid Intelligence and Response to Cognitive Training. Brain Topogr. 30, 502–520.

Seitzman, B.A., Abell, M., Bartley, S.C., Erickson, M.A., Bolbecker, A.R., Hetrick, W.P. (2017). Cognitive manipulation of brain electric microstates. Neuroimage 146, 533–543.

Simon, N.M., Otto, M.W., Wisniewski, S.R., Fossey, M., Sagduyu, K., Frank, E., et al. (2004). Anxiety disorder comorbidity in bipolar disorder patients: Data from the first 500 participants in the Systematic Treatment Enhancement Program for Bipolar Disorder (STEP-BD). Am. J. Psychiatry 161, 2222–2229.

Simon, N.M., Zalta, A.K., Otto, M.W., Ostacher, M.J., Fischmann, D., Chow, C.W., et al. (2007). The association of comorbid anxiety disorders with suicide attempts and suicidal ideation in outpatients with bipolar disorder. J. Psychiatr. Res. 41, 255–264.

Spielberger, C.D., Gorsuch, R.L., Lushene, R.E., (1970). Manual for the State-Trait Anxiety Inventory.

Strakowski, S.M., Adler, C.M., Almeida, J., Altshuler, L.L., Blumberg, H.P., Chang, K.D., et al. (2012) The functional neuroanatomy of bipolar disorder: A consensus model. Bipolar Disord. 14, 313–325.

Strik, W.K., Dierks, T., Becker, T., Lehmann, D., (1995). Larger topographical variance and decreased duration of brain electric microstates in depression. J. Neural Transmission, 99, 213–222.

Sverak, T., Albrechtova, L., Lamos, M., Rektorova, I., Ustohal, L., (2018). Intensive repetitive transcranial magnetic stimulation changes EEG microstates in schizophrenia: A pilot study. Schizophr. Res. 193, 451–452.

Tomescu, M.I., Rihs, T.A., Becker, R., Britz, J., Custo, A., Grouiller F., et al. (2014). Deviant dynamics of EEG resting state pattern in 22q11.2 deletion syndrome adolescents: A vulnerability marker of schizophrenia? Schizophr. Res. 157, 175–181.

Tomescu, M.I., Rihs, T.A., Roinishvili, M, Karahanoglu, F.I., Schneider, M., Menghetti, S., et al. (2015). Schizophrenia patients and 22q11.2 deletion syndrome adolescents at risk express the same deviant patterns of resting state EEG microstates: A candidate endophenotype of schizophrenia. Schizophr. Res. Cogn. 2, 159–165.

Van De Ville, D., Britz, J., Michel, C.M., (2010). EEG microstate sequences in healthy humans at rest reveal scale-free dynamics. Proc. Natl. Acad. Sci. 107, 18179–18184.

Vargas, C., López-Jaramillo, C., Vieta, E., (2013). A systematic literature review of resting state network-functional MRI in bipolar disorder. J. Affective Disord. 150, 727–735.

Vieta, E., Salagre, E., Grande, I., Carvalho, A.F., Fernandes, B.S., Berk, M., et al. (2018). Early intervention in Bipolar disorder. Am. J. Psychiatry 175, 411–426.

Wang, Y., Zhong, S., Jia, Y., Sun, Y., Wang, B., Liu, T., et al. (2016). Disrupted resting-state functional connectivity in nonmedicated bipolar disorder. Radiology 280, 529–536.

Wiedemann, G., Stevens, A., Pauli, P., Dengler, W., (1998). Decreased duration and altered topography of electroencephalographic microstates in patients with panic disorder. Psychiatry Res. Neuroimaging 84, 37–48.

Williams, J.B.W. and Kobak K.A., (2008). Development and reliability of a structured interview guide for the Montgomery Asberg depression rating scale (sigma). Br. J. Psychiatry 192, 52–58.

Yin, Z., Chang, M., Wei, S., Jiang, X., Zhou, Y., Cui, L., et al. (2018). Decreased functional connectivity in insular subregions in depressive episodes of bipolar disorder and major depressive disorder. Front. Neurosci. 12(NOV).

Young R.C., Biggs, J.T., Ziegler, V.E., Meyer, D.A., (1978). A rating scale for mania: Reliability, validity and sensitivity. Br. J. Psychiatry 133, 429–435.

